# A defined bacterial consortium and spatial transcriptomics highlight the complex interaction between *Campylobacter jejuni* and the murine intestine

**DOI:** 10.1101/2025.09.23.678081

**Authors:** Siddhi Chitre, Raad Z. Gharaibeh, Christian Jobin

## Abstract

The intestinal microbiota influences host susceptibility to *Campylobacter jejuni* (*C. jejuni*) infection. However, the interaction between specific intestinal bacteria and the *C. jejuni-*mediated host response is unclear. We established a defined consortium of bacteria to delineate *C. jejuni*-induced host responses. Three groups of germ-free (GF) *Il10*^-/-^ mice were used in this study: 1) mice colonized with a defined consortium of 13 bacterial isolates (C13) representing the four most prominent phyla in the mouse gut. 2) C13 plus *C. jejuni* 81-176 and 3) GF alone. The C13 + *C. jejuni* group induced significant intestinal inflammation and inflammatory mRNA gene expression compared to mice colonized with C13. 16S rRNA gene sequencing revealed an increased relative abundance of *Escherichia* and *Paraclostridium* in C13 + *C. jejuni*. Fluorescence *in situ* hybridization (FISH) and RNAscope integrated with spatial transcriptomics provided a high-resolution map of infection-induced gene expression, revealing localized immune responses and epithelial remodeling in defined colonic regions. Region-specific analysis further demonstrated that tissue-associated *C. jejuni* differentially modulates host gene expression compared to tissue-associated *Enterobacteriaceae*. Collectively, these findings demonstrate the potential of defined microbial consortia and spatially resolved transcriptomics to dissect the complex interplay between host, microbiota, and pathogens during enteric infection.

## Introduction

The gut microbiota comprises a rich and diverse community of microorganisms that co-exist in the gastrointestinal tract and is a prerequisite for maintaining gut homeostasis (1). These gut microbes participate in critical nutrient metabolism, immune system development, and protection against invading pathogens (2) including *Campylobacter jejuni* (*C. jejuni*) (3). This bacterium is a major cause of bacterial gastroenteritis worldwide and is associated with both acute illness and long-term complications, including Guillain–Barré syndrome and inflammatory bowel disease (IBD) (4). *C. jejuni* is an enteric pathogen that possesses several virulence factors, including cytolethal-distending toxin (CDT) virulence gene exhibiting DNA-damaging ability in the host (5,6). *C. jejuni* is commonly detected in the stool samples of pediatric patients with IBD (7). Interestingly, disease flare-ups and emergency room visits were higher for IBD patients infected with *C. jejuni* CDT-positive strains (8). In addition, *C. jejuni* infection favors the development of neoplastic lesions and metastasis in pre-clinical models, phenotypes dependent on the presence of CDT (9,10). We previously demonstrated that *C. jejuni* infection triggered rapid and severe colitis in germ-free *Il10^-/-^*mice (11,12) but not when these mice were colonized with microbiota before infection (3). However, studies involving conventional specific pathogen-free (SPF) mice remain difficult to interpret due to their complex and undefined microbial communities. In this study, we established a novel consortium of 13 bacteria (C13) representing the dominant bacterial phyla (13) in the mouse gut as a tractable system to study host response to *C. jejuni* 81-176 infection in GF *Il10^-/-^* mice. We observed that *C. jejuni* infection of C13-colonized mice triggered host inflammatory gene expression and increased *E. coli* colonization in the colons compared to C13 alone. We combined microbial and host analyses, including 16S rRNA gene sequencing, bacterial and host RNA-seq, histology, and qPCR. To gain spatial insights, we used fluorescence in situ hybridization (FISH) and RNAscope to localize bacterial and host transcripts, followed by spatial transcriptomics to examine region-specific gene expression patterns.

We demonstrate that the use of a bacterial consortium replicates all the major classic campylobacteriosis hallmarks, including increased inflammatory gene expression, neutrophil infiltration and *C. jejuni* tissue invasion. Spatial transcriptomic analysis showed that host responses to microbial presence are highly specific, emphasizing the critical role of local tissue microenvironments in shaping epithelial and immune dynamics to bacterial colonization.

## Methods

### Mice Experiments

All animal experiments were approved by the Institutional Animal Care and Use Committee (IACUC) at the University of Florida (UF) and performed at UF Animal Care Facilities (IACUC Protocol #202200000637). Colonies of germ-free mice were bred and maintained in isolators by the UF Animal Care Services Germfree Division. Mixed gender, 6-10 week old germ-free (GF *Il10^-^*^/-^) 129/SvEv mice were transferred from breeding isolators and placed into the Techniplast ISOcage P Bioexclusion system to allow for microbial manipulation (14,15). Cages were supplied with autoclaved food and water. Mice were randomly assigned to either GF group, thirteen-member consortium (C13) group or thirteen-member consortium with *C. jejuni* 81-176 (C13 + *C. jejuni*) group. Freshly passed stools were collected on day 6 using autoclaved forceps into sterile Eppendorf tubes and flash-frozen in liquid nitrogen and stored at -80°C. Colons were harvested and flushed with 1X PBS. One half of the proximal and distal colon snips were collected and flash-frozen in liquid nitrogen and stored at -80°C. The other half of the colon was swiss-rolled and fixed in 10% formalin overnight at room temperature (RT) and the next day transferred to 70% ethanol for processing and paraffin embedding.

### Histological assessment

The sections were stained with hematoxylin and eosin and the slides scanned using an Olympus VS200 microscope (UF molecular pathology core). Two investigators blinded to the treatment allocations scored inflammation based on the histology of the tissue. Briefly, intestinal inflammation was scored on a scale of 0-4 by the degree of lamina propria immune cell infiltration, goblet cell depletion, crypt hyperplasia, crypt abscess, and architecture distortion and were calculated as average of proximal, mid and distal colon regions as described previously (10).

### Bacterial culture

Members of the consortium were individually purchased from DSMZ (German Collection of Microorganisms and Cell Cultures GmbH; Braunschweig, Germany) representing four major bacterial phyla. *Bacteroides acidifaciens* (DSM 100502), *Bacteroides caecimuris* (DSM 26085), *Blautia caecimuris* (DSM 29492), *Thomasclavelia ramosa* (DSM 29357), *Parabacteroides goldsteinii* (DSM 29187), *Akkermansia muciniphila* (DSM 26127) and *Paraclostridium bifermentans* (DSM 29423). The bacterial strains *Parabacteroides distasonis* and *Bacteroides uniformis* were isolated by the Jobin’s laboratory from the stools of healthy mice. All the bacterial strains described above were grown from a glycerol stock in MEGA media (16) for 3 days under anaerobic conditions. *Escherichia coli* (DSM 28618) was grown aerobically in Luria Bertani broth. *Staphylococcus xylosus* (DSM 28566) was grown in tryptic soy broth aerobically. *Enterococcus hirae* (DSM 28619) was grown in tryptic soy broth and *Ligilactobacillus murinus* (DSM 28683) was grown in MRS medium microaerophically created by BD GasPak Gas generating systems. *Campylobacter jejuni* 81-176 was grown in Campylobacter Selective Medium (Fisher Cat # R01274) for 48 hours at 37°C using the BD GasPak EZ Campy pouch generating systems (Fisher Scientific Cat #B260685). CFU estimation was done using nanodrop for culture OD. Each isolate was pooled using equal CFU amounts to reach a total 1 x 10^6^ CFU/mouse dosage in 200 µL.

### Preparation of ISOcage P Bioexclusion system

Germ-free mice were maintained under sterile conditions using the ISOcage P Bioexclusion system. UF ACS staff transferred germ-free mice in a transfer disk to the biosafety cabinet. The mice were then carefully transferred using sterilized forceps to the pre-sterilized cages filled with autoclaved water and food. Cages and other materials were sterilized with Exspor (chlorine-dioxide sterilant activator and base) as described previously (17). For the stool collection, beakers, pre-labeled tubes, and forceps were autoclaved one day before collection. Post autoclave, the bag was immediately sealed with clear tape. On the day of stool collection, the same procedure was followed for Exspor sterilization for the cages, biosafety cabinet, and the autoclaved materials for stool collection. Freshly passed stools were collected in the tubes and were flash-frozen in liquid nitrogen and stored at -80°C until used for analysis.

### Fecal DNA Extraction, 16S rRNA Gene Sequencing and Analysis

DNA extraction was performed as described previously (18). One-half stool pellet per mouse was used for fecal DNA extraction, and samples were aliquoted in a 96-well PowerBeard Pro Plate (QIAGEN). Samples were lysed using Tissue Lyser II (QIAGEN) and then processed using QiAcube HT as described (19).

Following total fecal DNA extraction, the 16S rRNA V1-V3 hypervariable region was amplified using barcoded primer pairs 27F (5’-AGAGTTTGATCCTGGCTCAG-3’) and 534R (5’-ATTACCGCGGCTGCTGG-3’) with universal Illumina paired-end adapter sequences. PCR products were purified, quantified, and pooled as described previously and sequenced in a single run of Illumina MiSeq (20). Demultiplexed reads were fed to the DADA2 (21) pipeline for primer sequence removal, quality filtering, correction of Illumina amplicon sequencing errors and dereplication followed by amplicon sequence variants (ASVs) generation and chimera removal. Taxonomic classification was then performed using DADA2 assignTaxonom and addSpecies functions utilizing an in-house database composed of SILVA reference dataset (v. 138.1), GTDB (v. bac120_arc122_ssu_r89) and RefSeq-RDP16S (v. 3). We then removed any ASV that was classified as non-bacterial and all singleton ASVs. This resulted in a total of 1,042,049 reads, with an average of 69,470 reads per sample (min= 58,880; max= 81,295).

Differential taxa abundance analysis was performed using MaAsLin2 (22) R package with group set as fix effect and cage set as a random effect. For barplot representation of the proportional relative abundance of taxa, we agglomerated the ASVs at the species level using phyloseq (23) package and then used R ggplot package for plotting.

### Stool and colonic tissue RNA extraction

Total RNA was isolated from frozen mouse stool day 6 samples as described (24). Briefly, lysis and disruption steps were performed with 1:1 mix of 1mm acid-washed glass beads and 0.1 mm zircona beads using a bead beater for efficient lysis and disruption; Precellys24 (Bertin Instruments Cat # EQ03119-200- RD000.0). RNA purification was further performed according to the manufacturer’s protocol using mirVana miRNA isolation kit, with phenol (Fisher Scientific AM1560). RNA eluted was treated with Turbo-DNA-free kit to remove any DNA impurities (Fisher Cat # AM1907). Colonic tissue RNA was extracted using TRIzol and a bead beater as previously described (25). RNA concentration was quantified using a NanoDrop 2000.

### RNA sequencing and analysis

Quality control, rRNA depletion and cDNA library preparation were performed by the University of Florida’s Interdisciplinary Center for Biotechnology Research (ICBR) Gene Expression Core (https://biotech.ufl.edu/gene-expression-genotyping/ RRID:SCR_019145). Briefly, RNA was quantified using the QUBIT fluorescent method (Fisher Biotech), and sample quality was assessed using the Agilent 2100 Bioanalyzer (Agilent Technologies, Inc.). Ribosomal RNAs were removed from total RNA using Illumina Ribo-Zero Plus rRNA Depletion Kit following the manufacturer’s protocol and eluted into 5 µl EB buffer. Then, the RNA-seq library was processed using NEBNext® Ultra™ Directional RNA Library Prep Kit for Illumina (NEB, USA) following the manufacturer’s recommendations. The libraries were pooled at equal molar concentrations and sequenced by NovaSeq X 10B flow cell with 2x150 cycles run. Sequencing was performed at the ICBR NextGen Sequencing (https://biotech.ufl.edu/next-gen-dna/, RRID:SCR_019152).

For RNA-seq analysis, reads were quality trimmed at Q20 (sequencing quality score of 20) and filtered to remove sequencing adaptors using Trimmomatic (26) (v.0.39). Alignment and quantification were done by STAR (27) (v.2.7.11b) using mouse reference genome (GRCm39) with its corresponding annotation file obtained from the National Center for Biotechnology Information (NCBI). An average of 14,367,266 reads per sample (min= 645,786; max= 40,516,378) were uniquely mapped to GRCm39. Gene counts were then imported into edgeR (28) for normalization and differential expression analysis. We considered a gene differentially expressed if its edgeR *P*_adj_-value is less than 0.05. Principal Component Analysis (PCA) was done using R prcomp function using the output of R DESeq2 rlog (regularized log transformation) function, which produces a matrix of log transformed and normalized counts, with the option blind set to true. We tested for differences in samples clustering using gls (Generalized Least Squares model) function in R package nlme.

The pathway analysis was generated using QIAGEN IPA (QIAGEN Inc., https://digitalinsights.qiagen.com/IPA) (29) software from UF-Health Cancer Center ICBR Bioinformatics core (RRID:SCR_019120).

For Bacterial RNA-seq analysis, trimmed and adaptor free RNA-seq reads from above were fed into Kraken2 (30) and classified using the nt database v.k2_nt_20231129 (a large collection of GenBank, RefSeq, TPA and PDB sequences) obtained from https://benlangmead.github.io/aws-indexes/k2. We then collected reads that were classified as bacteria (average of 28,230,937 bacterial reads per sample; min= 7,762,062, max= 37,199,160) and used them for subsequent bacterial RNA-seq analysis, which guarantee that reads from host and other organisms do not introduce any contamination into our analysis. Those reads were then aligned to a reference composed of the genomes of our bacterial consortium (obtained from ftp://ftp.ncbi.nlm.nih.gov/genomes) using bowtie2 (31) aligner (average of 91% of input reads successfully aligned; min=81%; max = 96%). Gene expression was quantified using featureCounts (32) from subread package. Counts were imported to edgeR and analyzed as described under Mouse RNA-seq analysis above. KEGG Orthology (KOs) abundances were generated using fmh-funprofiler (33) from the same reads used for bowtie2 alignment above, then pathway analysis was conducted through GAGE (34) using KEGG pathways. We considered a pathway significant if its GAGE *P*_adj_-value is less than 0.05.

### Immunohistochemistry (IHC)

Neutrophils in colon tissues were detected using immunohistochemistry for the marker Ly-6G/Ly-6C (Gr- 1; Clone RB6-8C5) as described previously (35). Colonic tissues were deparaffinized, blocked, and incubated overnight with Ly-6G/Ly-6C (1:100). The next day, tissues were incubated with anti-rabbit biotinylated (NBP1-75414 -1:500), ABC (Vectastain ABC Elite kit, Vector Laboratories), DAB (Dako), and hematoxylin. The images were captured using an Olympus VS200 slide scanner (Evident Scientific) at 40X magnification to assess brown staining. The signals were quantified on ImageJ, where five fields of vision were counted.

### Fluoresence in situ hybridization (FISH)

FISH assay was performed as previously described (11). Briefly, tissue sections from all groups were deparaffinized, hybridized, washed, and mounted in DAPI medium and imaged using a Leica DM6000B upright microscope. Cy3- tagged 5’AGCTAACCACACCTTATACCG3’ targetting *C. jejuni* 23S rRNA (36) was used to detect *C. jejuni* localization in the colons.

### RNAscope

The RNAscope 2.5HD Duplex Detection kit (Cat no.322430), RNAscope probe for *Enterobacteriaceae*- 16S mRNA (Cat no. 509831), and mouse tumor necrosis factor (*Tnfa*) mRNA (Cat no.311081-C2) were purchased from Advanced Cell Diagnostics (ACD-Biotechne). RNA scope was performed according to the manufacturer’s instructions. The slides were captured using an Olympus VS200 slide scanner (Evident Scientific) at 40X magnification to assess positive cells. The signals were quantified on ImageJ, where five fields of vision were counted. Qualitatively, fewer positive signals were observed in low regions compared to high regions of the mid-colon, where a denser distribution was evident, providing the basis for their classification as ‘low’ and ‘high’ regions. We also utilized slide scanning service (Olympus VS200) operated by UF Molecular Pathology Core (RRID:SCR_016601).

### 10X Visium HD Spatial Transcriptomics

Formalin-fixed, paraffin-embedded (FFPE) tissue blocks were used. Prior to sectioning, RNA quality was assessed by extracting RNA from test sections and calculating the DV200 value (>30% required for optimal spatial transcriptomics). FFPE blocks were faced at 10 μm to expose the tissue, then sectioned at 5 μm thickness using a microtome. Sections were dried at 42°C for 3 hours and stored overnight in a desiccator at room temperature. Spatial transcriptomics was performed by UF ICBR (RRID:SCR_019145), according to the manufacturer’s protocol. Spatially barcoded cDNA was extracted and used for library construction following the Visium for FFPE library prep protocol. Libraries were quantified using a Bioanalyzer (Agilent Technologies) and sequenced on an Illumina NovaSeq 6000 platform using 150 bp paired-end reads, targeting 50,000 read pairs per capture spot resulting in more than 48 million paired-end reads per sample.

### Spatial Transcriptomics Data Processing and Analysis

FASTQ files were processed using Space Ranger (v3.1.1) using mouse reference genome (mm10) resulting in more than 99% of reads mapped to probe set, more than 97% of reads mapped confidently to probe set, more than 97% of reads mapped confidently to the filtered probe set, more than 92% of barcodes are valid, more than 99% of UMIs are valid, more than 96% of reads in squares under tissue and less than 1.2% of UMIs are estimated from Genomic DNA. The resulting Space Ranger outputs were imported to Seurat R package (v5.2.1) for analysis following Seurat spatial transcriptomics analysis workflow: https://satijalab.org/seurat/articles/spatial_vignette.html. Briefly, we included spots with a percentage of mitochondrial genes less than 20% and have counts greater than 100. Data were normalized and variance stabilized using Seurat’s SCTransform. For analysis involving different slices, Harmony (v.1.2.3) (37) was used for batch effect correction, followed by PrepSCTFindMarkers. For region of interest analysis, the selected areas contained roughly the same number of bins. We used single-cell RNA sequencing data from Mayassi et al. (38) as a reference for cell-type deconvolution in our spatial transcriptomics data using spacexr v.2.2.1 (39).

### RT-qPCR for *E. coli* gene transcripts

500 ng of RNA was reverse transcribed using the iScript cDNA Synthesis Kit (Bio-Rad 170-8891). qPCR was performed using a Bio-RAD CFX384 Real-Time PCR system using primers described in Table 1 (*E. coli* virulent transcripts). Amplification was performed using the following conditions: initial denaturation at 95°C for 3 minutes, followed by 39 cycles of 95°C for 10 seconds, 56°C for 10 seconds, and 72°C for 20 seconds. The melting curve was performed by increasing the temperature from 65°C to 95°C in 0.5°C increments, with a 5-second hold at each step. The universal bacterial 16S ribosomal gene was used as a control. Fold change was calculated using the 2^-ΔΔCt^ method (40) using SYBR green dye (Applied Biosystems).

**Table 1:**
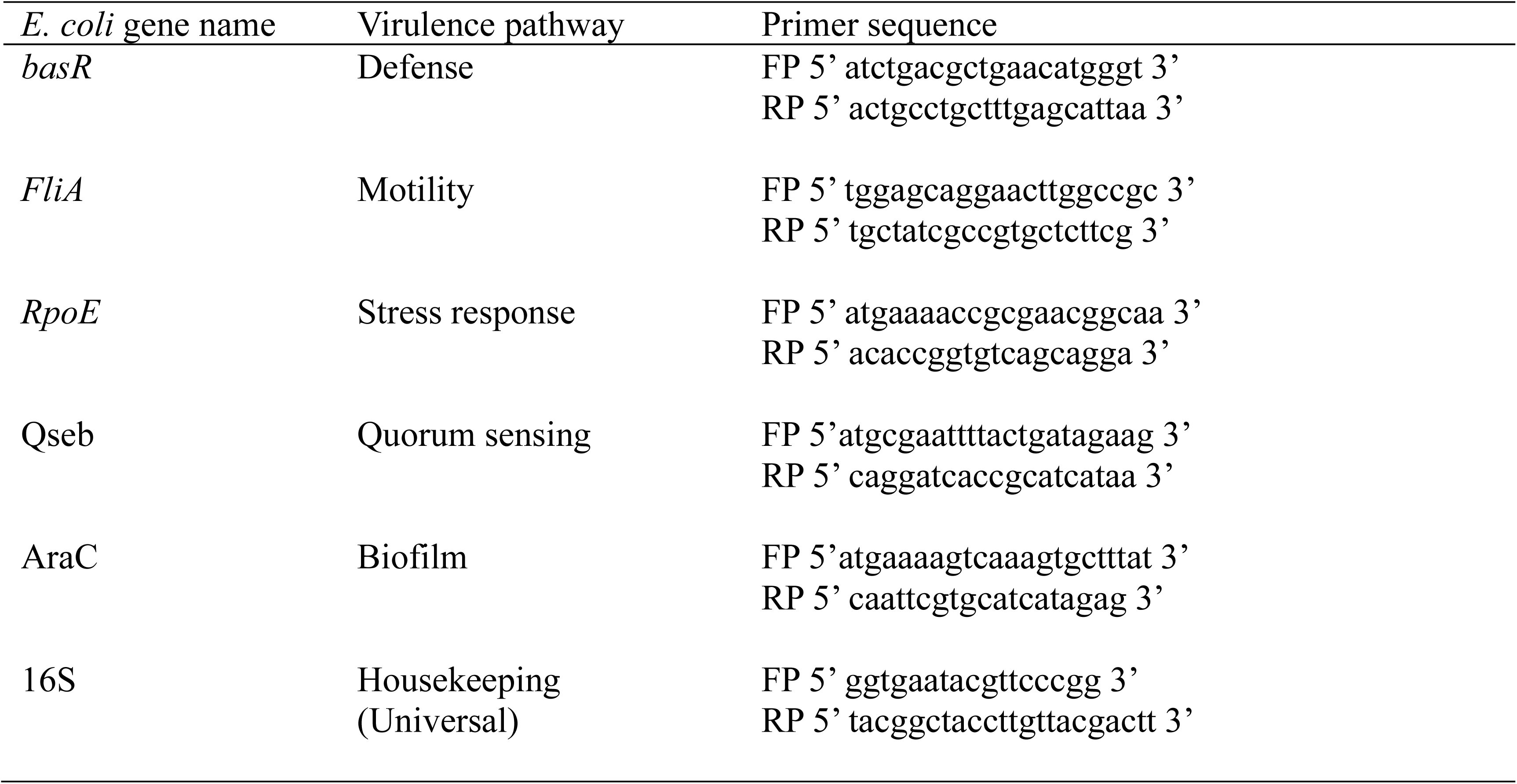
Primers used for RT-qPCR for *E. coli* gene transcripts.

### RT-qPCR for mammalian gene transcripts

500 ng of RNA was reverse transcribed and qPCR was performed using a Bio-RAD CFX384 Real-Time PCR system using primers described in Table 2 (host gene transcripts). Amplification was performed using the following conditions: initial denaturation at 95°C for 3 minutes, followed by 39 cycles of 95°C for 10 seconds, 56°C for 10 seconds, and 72°C for 10 seconds. The melting curve was performed by increasing the temperature from 65°C to 95°C in 0.5°C increments, with a 5-second hold at each step. *Gapdh* was used as a housekeeping control. Fold change was calculated using the 2^-ΔΔCt^ method (81) using SYBR green dye (Applied Biosystems).

**Table 2:**
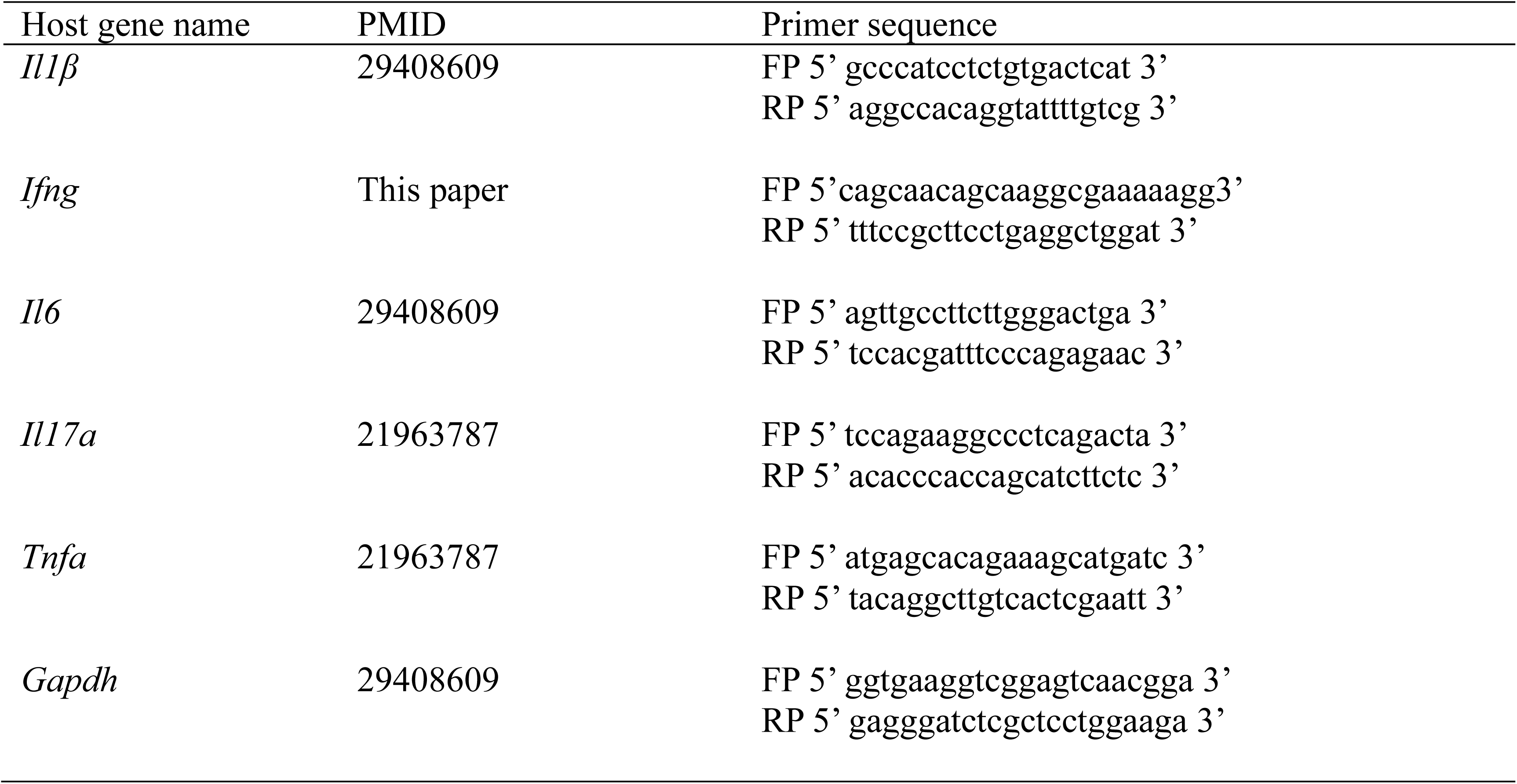
Primers used for RT-qPCR for host inflammatory gene transcripts.

### Statistical tests

We used R’s two-sample Wilcoxon test (also known as Mann-Whitney test) and for tests involving more than two comparisons, we corrected the p-values for multiple testing using R’s p.adjust function with method set to BH (Benjamini and Hochberg) (41).

## Results

### *C. jejuni* infection drives severe inflammation and increased abundance of *E. coli* and *P. bifermentans* in the colon

To establish a tractable system to study host response to *C. jejuni* infection, we assembled a consortium of 13 bacterial species representing 4 major phyla present in the intestine. Since susceptibility to *C. jejuni* infection is influenced by the microbiota (3), we first examined the impact of the interaction between C13 and *C. jejuni* on host response using *Il10^-/-^* germ-free (GF) mice, a well-established model of campylobacteriosis (12,42). GF *Il10^-/-^* mice were divided into three groups: GF, C13 and C13 containing *C. jejuni*. Each group was colonized with their respective bacterial community or no gavage (single oral gavage, 10^6^ CFU/mouse) and housed in an ISOcage P Bioexclusion cage system. Six days post-oral gavage, stool and distal colon samples were collected from all the mice (Fig.1A). To determine engraftment of the consortium and impact of *C. jejuni* on C13, we profiled bacterial composition using 16S rRNA gene sequencing. We observed effective colonization of each member of the consortium in both C13 and C13 +

**Figure 1.**
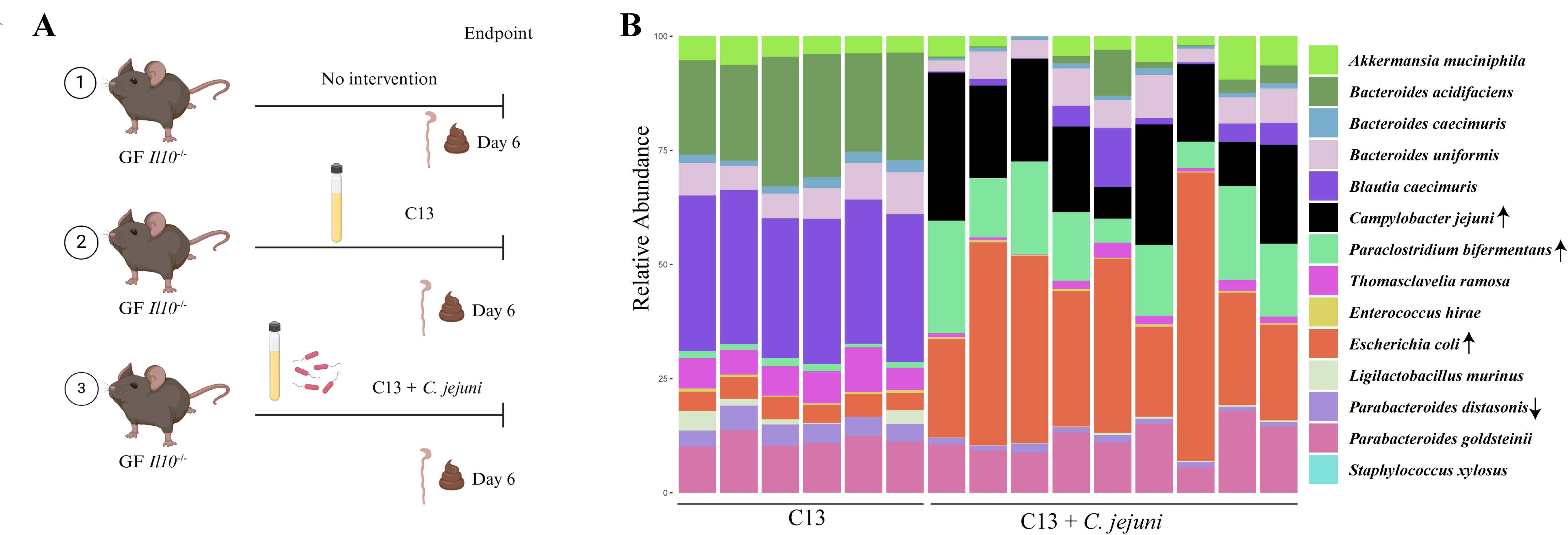
C13 + *C. jejuni* infection increased the relative abundance of *Escherichia* and *Paraclostridium* (A) Experimental workflow for investigating host-bacterial communication in C13 and C13 + *C. jejuni* infection. (B) Barplot shows the relative abundance of the bacterial consortium present in the C13 and C13 + *C. jejuni* fecal samples as determined by 16S rRNA gene sequencing, with each bar representing a single mouse. Up arrow indicates significant enrichment (*P*_adj_ < 0.05) in C13 + *C. jejuni*, and the down arrow indicates significant depletion (*P*_adj_ < 0.05) in C13 + *C. jejuni*.

*C. jejuni*-colonized *Il10^-/-^* germ-free mice (Fig.1B). We observed significant expansion of *Escherichia coli* and *Paraclostridium bifermentans* and contraction of *Parabacteroides distasonis* relative abundance in the C13 + *C. jejuni* group compared to C13 alone. Consistent with our findings, previous 16S rRNA sequencing studies have reported an increased abundance of *Enterobacteriaceae* following *C. jejuni* infection of antibiotic-treated mice (43), supporting the role of *C. jejuni* in driving microbial community shifts that may contribute to inflammation.

*C. jejuni* infection triggers rapid colitis in GF *Il10^-/-^* mice but remains healthy when mice are colonized with microbiota (3). Therefore, we next compared the severity of inflammation between C13 and C13 + C. *jejuni* colonized mice. While *Il10^-/-^* mice colonized with C13 displayed minimal inflammatory response, the addition of *C. jejuni* to the consortium provoked significantly increased inflammation as quantified by histological score with increased immune cell infiltration, loss of goblet cells and crypt abscesses (Fig. 2A, B). Similarly, a significant upregulation of inflammatory genes such as *Ifng*, *Il1b, Il6*, *Tnfa*, and *Il17a* mRNA was observed in C13 + *C. jejuni*-infected mice compared to C13 mice (Fig. 2C-G). Thus, a defined- bacterial consortium allows the study of campylobacteriosis in *Il10^-/-^* mice.

**Figure 2.**
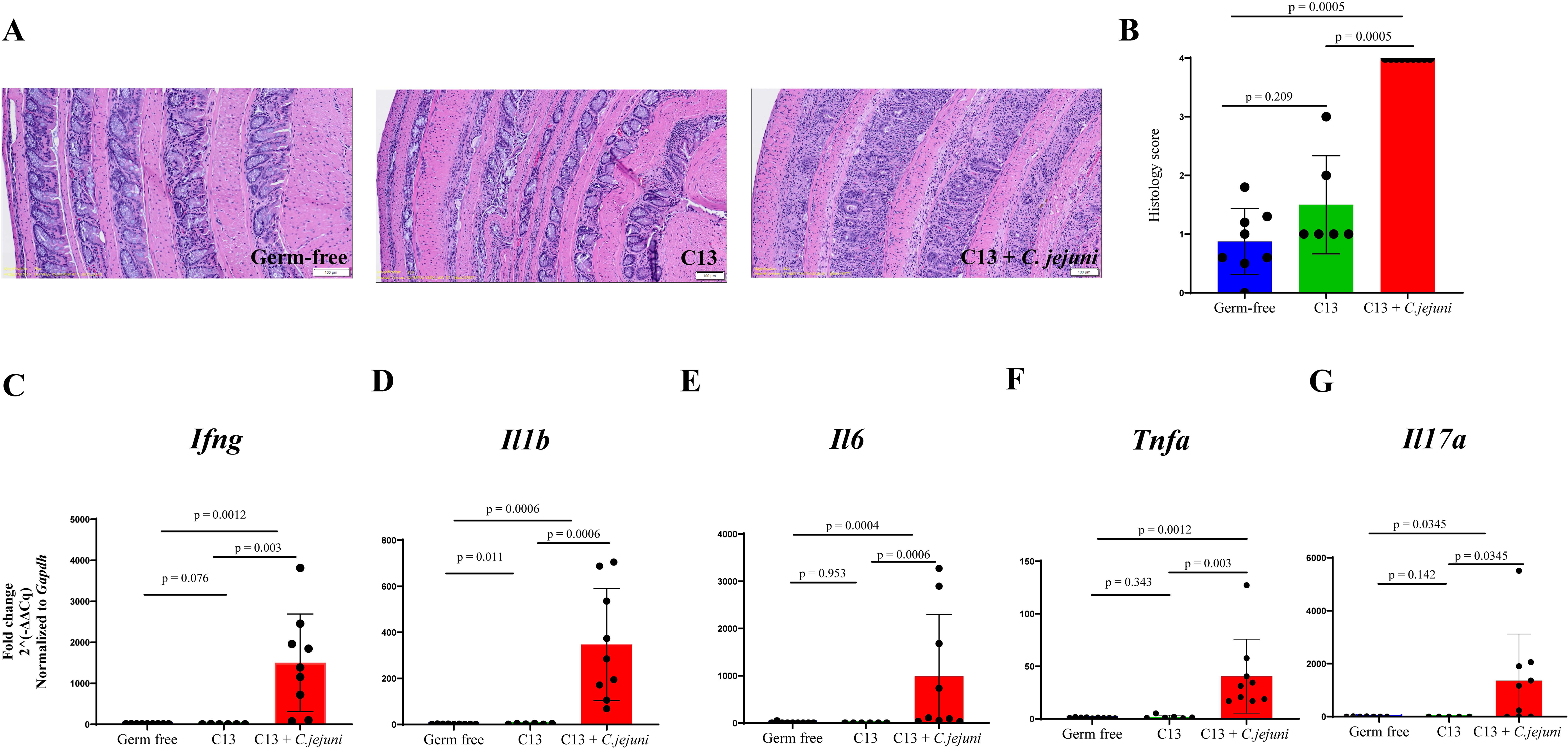
Severe inflammation observed in C13 + *C. jejuni* infected mice compared to C13 (A) H&E stained colonic swiss rolls of different experimental groups illustrating histopathological differences between groups. (B) Quantification of colon histological scores, comparing inflammation across experimental groups. (C-G) RT-qPCR analysis of inflammatory mouse gene transcript *Ifng, Il1b, Il6, Tnfα, and Il17a*. For histology scores and RT-qPCR analysis, *P*_adj_-values were calculated using R’s two-sample Wilcoxon test and corrected for multiple testing.

### C13 + *C. jejuni* infection modulates host and bacterial transcriptomic profile as compared to C13 colonized mice

To gain a better understanding of the changes in host physiology caused by *C. jejuni* infection, we performed bulk RNA-seq from stool specimens. This approach allowed us to study both gene expressions from exfoliated gut luminal epithelial cells as well as associated microbial gene transcripts. PCA showed that *C. jejuni* infection induced marked changes in the host transcriptional response (Fig. 3A). We observed 4,085 significant differentially expressed host genes, with 2,221 upregulated and 1,864 downregulated in C13 + *C. jejuni* colonized mice (Fig. 3B, Additional file 1: https://figshare.com/s/b84ac4a3c99882822195). We noted increased expression of *Tgm3* in C13 colonized mice, a gene implicated in mucosal barrier function, preventing pathogenic infections (44), and *Gsdmc2* and *Gsmdc3* genes which play a role in Lgr5^+^ stem cell maintenance and regeneration by regulating mitochondrial homeostasis (45). In addition, we observed that the *Slc7a11* gene, which plays a role in extracellular cysteine uptake and is essential for maintaining redox balance (46), is upregulated in C13 + *C. jejuni* mice compared to C13 colonized mice. We also noted increased chemokine *Cxcl5*, a chemoattractant for neutrophil infiltration (47) and *Duoxa2* expression, a gene necessary for activation of the dual oxidases DUOX2 implicated in anti-microbial host response (48). Using Qiagen Ingenuity pathway analysis (IPA), we found that different immune response pathways, including neutrophil degranulation pathway, pathogen-induced cytokine storm, leukocyte extravasation, and NOD1/2 pathway were significantly upregulated upon *C. jejuni* infection (Fig. 3C).

**Figure 3.**
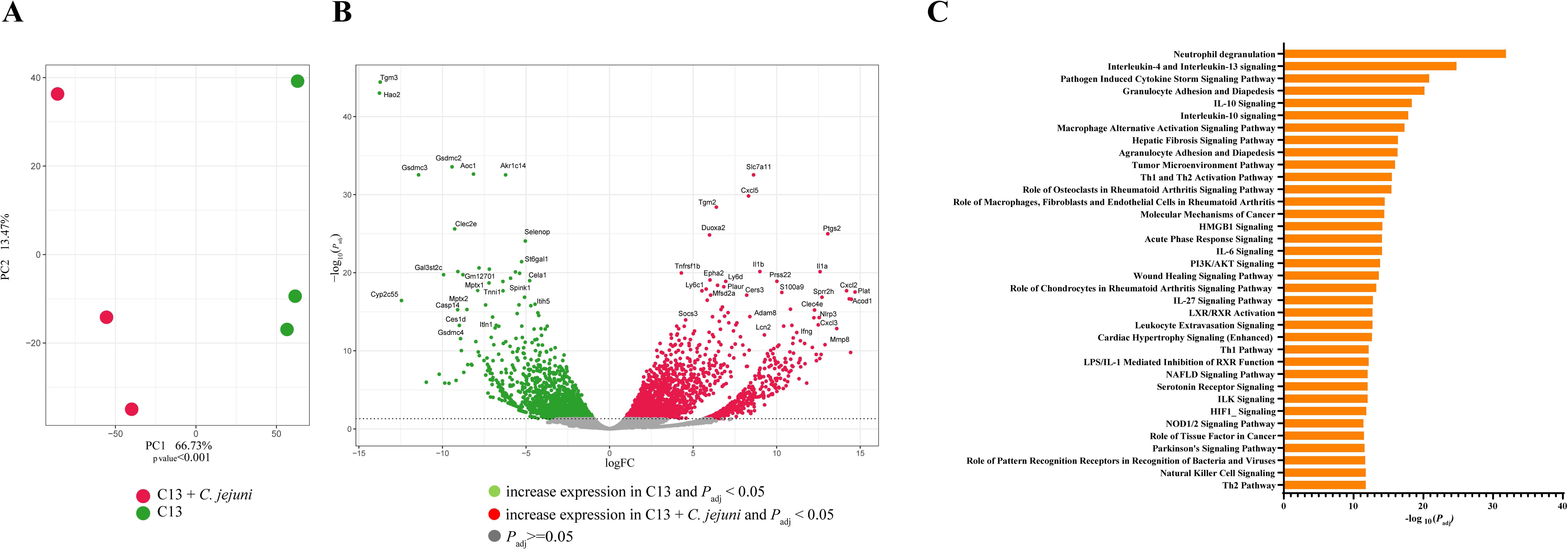
Transcriptome analysis reveals alterations in host gene expression following *C. jejuni* infection (A) PCA plot showing sample clustering based on their transcriptome in C13 and C13 + *C. jejuni* groups (p value < 0.01). Differences in sample clustering were tested using R’s gls. (B) Volcano plot of differentially expressed mouse genes highlighting the top genes significantly overexpressed in C13 and C13 + *C. jejuni* groups. Horizontal dotted line showing -log_10_(0.05). (C) Top host pathways significantly upregulated in C13 + *C. jejuni* mice as compared to C13. Qiagen IPA software was used for this analysis (*P*_adj_ value<0.05).

We next focused on the impact of *C. jejuni* infection on bacterial gene expression profiles. PCA showed distinct changes in the bacterial transcriptional profile following the addition of *C. jejuni* to the C13 (Fig. 4A). Volcano plot showed a plethora of differentially expressed genes between C13 and C13 + *C. jejuni* groups (Fig. 4B, Additional file 2: https://figshare.com/s/b84ac4a3c99882822195). As compared to the C13 group, we observed 8,754 downregulated and 7,327 upregulated bacterial genes following inclusion of *C. jejuni*. We observed a decrease in bacterial expression genes such as *Blautia* carbohydrate ABC transporter permease, *Ligilactobacillus* alpha glucoside-specific transporter, and *Bacteroides* RagB/ SusD outer membrane protein that play a role in nutrient uptake and in carbohydrate metabolism that aids in bacterial growth and survival (49,50). In contrast, we observed an increased expression of various bacterial genes, including *C. jejuni* PorA implicated in adaptation to change in the intestinal environment (51), *C. jejuni* and *P. bifermentans* peroxiredoxin (Prxs) and *C. jejuni* Dps family protein that confers protection from oxidative damage (52,53). Other notable altered genes included aspartate-ammonia lyase, which promotes *E. coli* growth under anaerobic conditions in the gut (54). These findings suggest that the presence of *C. jejuni* modified the bacterial transcriptomic profile in the gut to promote pathogen survival.

**Figure 4.**
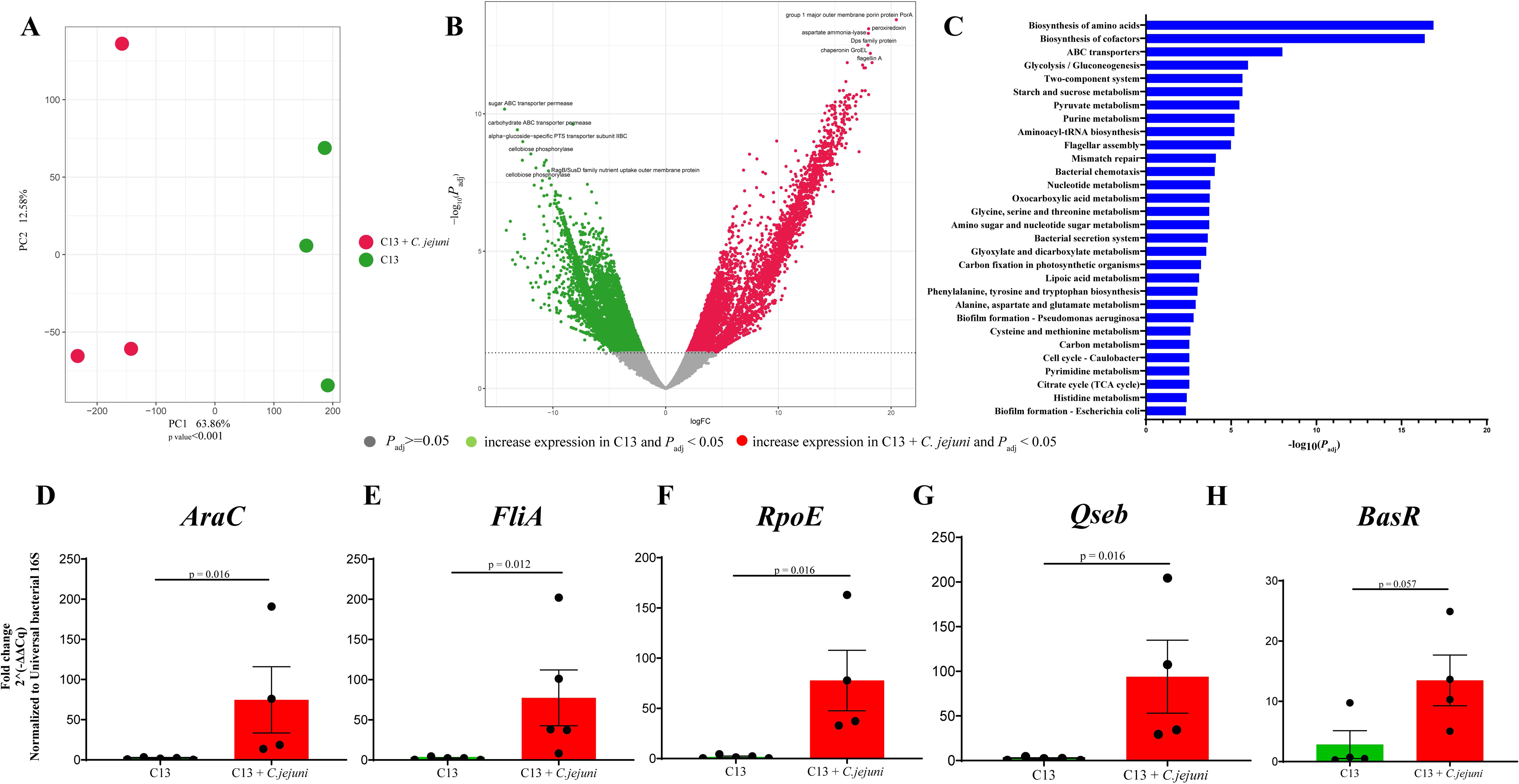
Pathway analysis reveals an increase in bacterial virulence pathways following *C. jejuni* infection (A) PCA plot showing sample clustering based on their bacterial transcriptome profiles in C13 and C13 + *C. jejuni* groups (p value < 0.01). Differences in sample clustering were tested using R’s gls. (B) Volcano plot of differentially expressed bacterial genes in C13 and C13 + *C. jejuni* (*P*_adj_ value <0.05). Horizontal dotted line showing -log_10_(0.05). (C) Top bacterial pathways significantly upregulated in C13 + *C. jejuni* group (GAGE *P*_adj_-value < 0.05). (D-H) RT-qPCR for *E. coli* virulence genes in stool samples of C13 and C13 + *C. jejuni* groups. P-values from R’s two-sample Wilcoxon test.

Using Kyoto Encyclopedia of Genes and Genomes (KEGG) pathway analysis, we observed that various bacterial metabolic pathways, such as the biosynthesis of amino acids, biosynthesis of co-factors, ABC transporters, purine, and pyruvate metabolism, were altered in C13 + *C. jejuni* mice compared to C13 (Fig. 4C). We observed that among significantly altered bacterial genes, 74.51% belonged to metabolic pathways, 17.65% to virulence pathways, and 7.84% to cell cycle and repair pathways in the C13 + *C. jejuni* colonized mice.

To validate these findings, we identified genes corresponding to virulent pathways present in *E. coli* such as biofilm, stress response, defense, motility, and quorum sensing. We selected a candidate gene for each pathway and normalized it to the universal bacterial 16S ribosomal gene using RT-qPCR. We observed a significant increase in *E. coli* AraC (biofilm), FliA (motility), RpoE (stress response) and Qseb (quorum sensing) transcripts in C13 + *C. jejuni* as compared to C13 (Fig. 4D-G). Although we observed an increase in *E. coli* BasR gene (defense), it was not statistically significant (*P*_adj_-value = 0.057, Fig. 4H). These data suggest that *C. jejuni* infection alters host inflammatory pathways and upregulates bacterial virulence pathways across the consortium.

### Presence of tissue invasive *C. jejuni* increased the abundance of *Enterobacteriaceae* in the colons of C13 + *C. jejuni* colonized mice

We previously observed significant expansion of *E. coli* in the stool samples of C13 + *C. jejuni*-infected mice (Fig. 1B). Thus, we next investigated the *C. jejuni* and *E. coli* distribution in the colon tissue. Fluorescence-in-situ hybridization (FISH) assay revealed *C. jejuni* infiltration in the mid colons of C13 + *C. jejuni*-infected mice compared to C13 mice (Fig. 5A). Interestingly, RNAscope analysis showed an increased abundance of *Tnfα* and *Enterobacteriaceae* mRNA transcripts in the C13 + *C. jejuni* group compared to C13 (Fig. 5B). Infiltration of *E. coli* following *C. jejuni* infection was also localized in the mid colon while *Tnfα* was dispersed throughout the colon (Fig. 5B). We previously demonstrated that neutrophils play a key role in mediating campylobacteriosis in *Il10^-/-^* mice (11,35), therefore we next sought to determine neutrophil infiltration in the colon of C13 + *C. jejuni* mice using immunohistochemistry (11,35). We observed significant neutrophil infiltration (Ly-6G^+^ marker) in the colons of C13 + *C. jejuni* compared to C13 (Fig. 5C). These findings demonstrate that the C13 is a tractable bacterial consortium enabling the study of host response to *C. jejuni* infection.

**Figure 5.**
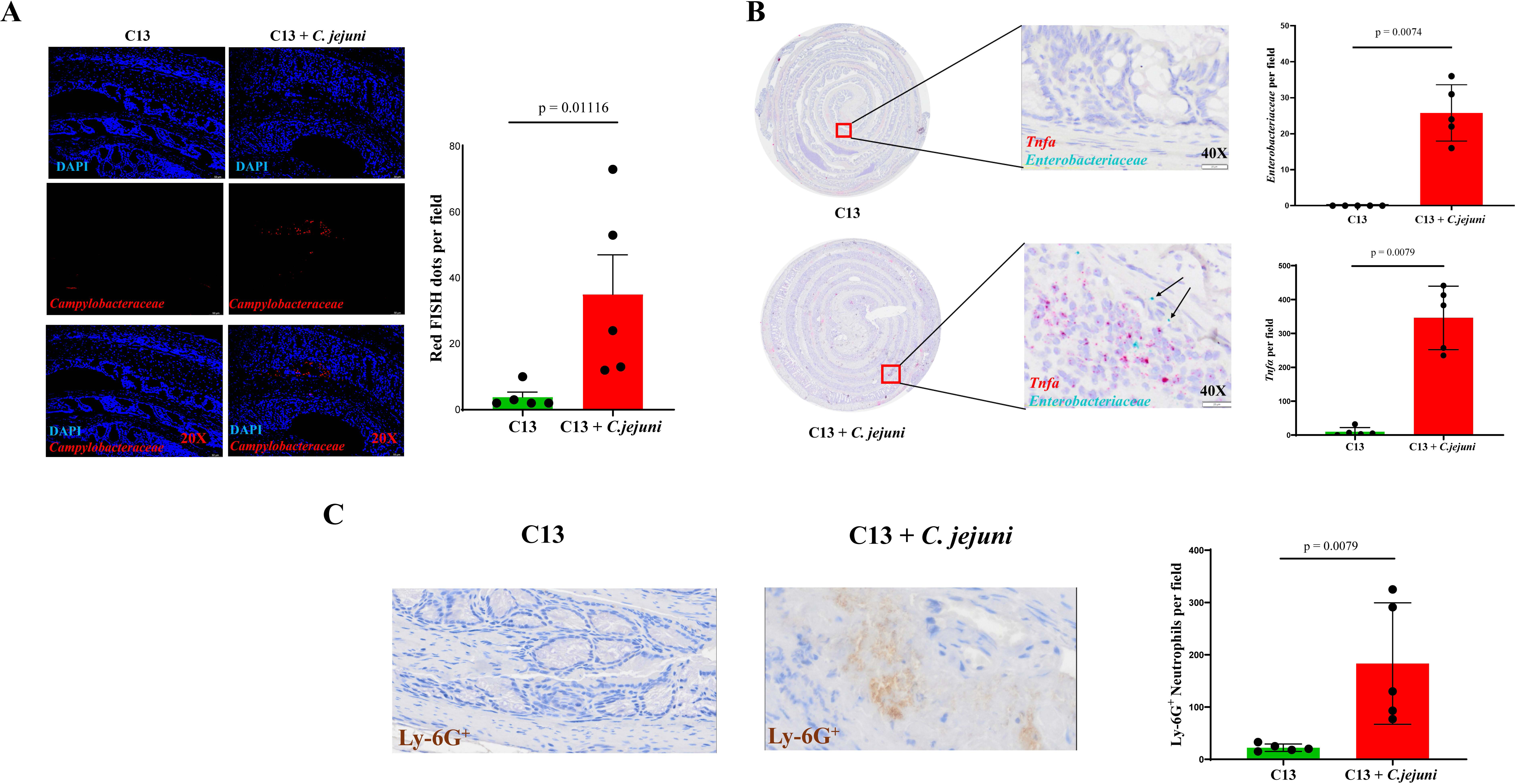
Increased *C. jejuni* and *Enterobacteriaceae* abundance accompanied with neutrophil infiltration in C13 + *C. jejuni* infected colons (A) Detection of *C. jejuni* as represented by red dots in colonic sections of colonized mice were detected using FISH. (B) Detection of *Enterobacteriaceae* and host *Tnfa* mRNA transcript expression using RNAscope. (C) Neutrophil detection by anti-Ly-6G antibody using IHC and quantification in the colons of C13 and C13 *+ C. jejuni* infected mice. p-values from R’s two-sample Wilcoxon test for FISH, RNAscope, and IHC assays.

### Spatial clustering reveals a distinct transcriptomic profile upon *C. jejuni* infection

The region-specific tissue colonization of *C. jejuni* and *Enterobacteriaceae* in the C13 + *C. jejuni-*infected mice prompted us to characterize the transcriptional reprogramming of these tissues. Clustering of our spatial transcriptomics revealed 19 (Clusters 0-18) clusters (Fig. 6 A) corresponding to distinct transcriptional signatures in the colonic tissue. We observed a similar clustering pattern between GF and C13-colonized mice, whereas the presence of *C. jejuni* affected several clusters (Fig. 6 A-C, Additional file 3: https://figshare.com/s/b84ac4a3c99882822195). For example, in the C13 + *C. jejuni* group, Clusters 2, 3, and 17 exhibited distinct gene expression patterns and spatial distributions throughout the colon, as visualized by spatial mapping (Fig. 6C, lower panel). Differential gene expression analysis identified cluster-specific marker genes for Clusters 2, 3, and 17, which were further characterized using IPA to assess enriched functional pathways.

**Figure 6.**
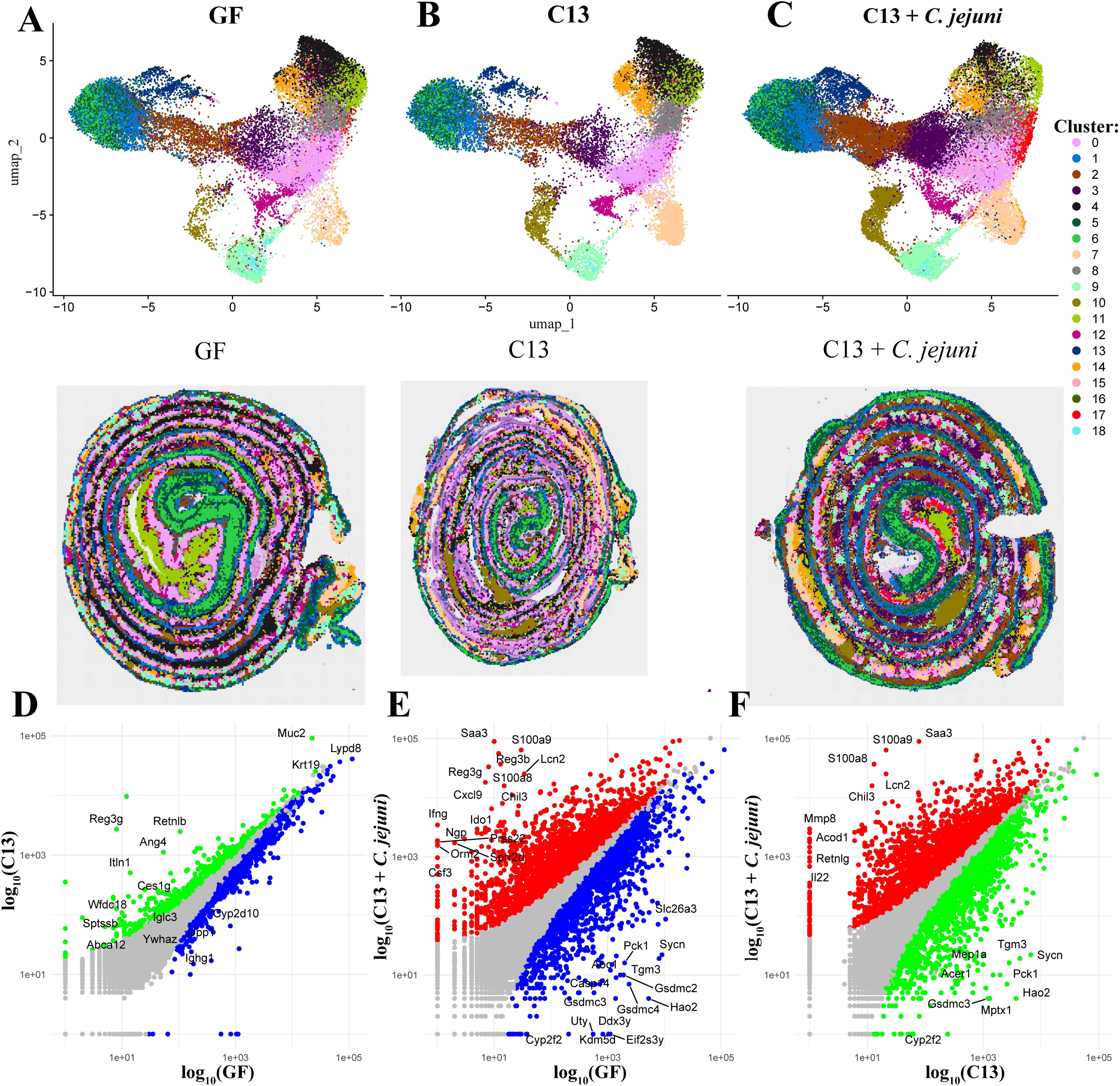
Spatial transcriptomics reveals distinct transcriptional clustering in the colons of GF, C13 and C13 *+ C. jejuni* mice (D-F) Scatter plot showing significantly differentially expressed genes in C13 vs GF, C13 *+ C. jejuni* vs GF and C13 *+ C. jejuni* vs C13. Each dot represents a gene; gray: *P*_adj_ ≥ 0.05, green: increased expression in C13 and *P*_adj_ < 0.05; blue: increased expression in GF and *P*_adj_ < 0.05; red: increased expression in C13 *+ C. jejuni* and *P*_adj_ < 0.05.

Cluster 2 showed significant upregulation of inflammatory genes associated with the S100 family signaling pathway, Th2 pathway, neutrophil degranulation, *Stat-3* and Interleukin signaling. Conversely, pathways involved in epithelial regulation and tumor suppression were downregulated, suggesting a localized pro-inflammatory environment (Supplemental Fig. S1A, Additional file 4: https://figshare.com/s/b84ac4a3c99882822195). Cluster 3 shared substantial overlaps with Cluster 2; to focus on distinct features, we selected pathways that were uniquely enriched in Cluster 3. These included pathways like Th1, PI3K/AKT, and NOD1/2 signaling pathways, suggesting coordinated activation of innate and adaptive immune responses in this cluster. Additionally, downregulation of lipid metabolism, LXR/RXR activation, and amino acid transport pathways pointed toward a metabolic shift (Supplemental Fig. S1B).

Cluster 17 demonstrated a distinct immunological and metabolic signature. Upregulated pathways included neutrophil extracellular trap formation, PTEN signaling, HMGB1, and glycolysis and gluconeogenesis, consistent with a stress-responsive environment. In contrast, downregulated pathways such as extracellular matrix organization, Rho family GTPase signaling, senescence, and phagosome formation implying reduced inflammation within this cluster (Supplemental Fig. S1C). Interestingly, Cluster 17 exhibited downregulation of pathways associated with pathogen-induced cytokine storm, contrasting with the upregulation observed in other clusters. Cluster 17 is predominantly localized to the distal colon, while *C. jejuni* abundance is higher in the mid colon, corresponding to clusters such as 2 and 3, which display increased inflammatory pathway activity. This spatial distribution highlights region-specific differences in gene expression during infection.

These results illustrate how host gene expressions are organized spatially in response to GF, C13, and C13 + *C. jejuni* infection. Overall, these data suggest that *C. jejuni* infection in the context of a defined consortia induces both widespread and localized transcriptional responses across the colonic tissue.

Furthermore, transcriptomic analysis of the entire colon revealed additional insight into overall gene expression changes across the different experimental groups. A total of 2,773 genes were significantly differentially expressed, of which 1,788 were downregulated and 985 were upregulated in C13 compared to the GF group (Fig. 6D). We compared our GF and C13 transcriptomic datasets to recently published spatial transcriptomic data from SPF mice (38). Interestingly, our C13 transcriptomic data overlaps with the published SPF mice dataset of Mayassi et al. (38), as evidenced by increased gene expression of microbe-driven host genes *Reg3g, Ang4, Itln1*, and *Wfdc18* in our C13 dataset, which play a role in host defense and mucosal homeostasis, compared to GF mice. The presence of *C. jejuni* in C13 drastically modified the transcriptomic landscape with upregulation of genes involved in antimicrobial defense (*Saa3, Reg3β, Reg3γ*), immune cell recruitment (*Ifnγ, S100a8, S100a9*), inflammation and barrier maintenance (*Prss22, Chil3*), tissue repair (*Il22, Mmp8*), neutrophil function (*Retnlg*), and immune metabolism (*Acod1*) (Fig. 6E-F). These overlapping responses suggest that C13 + *C. jejuni* maintains a distinct gene expression profile relative to both GF and C13 mice.

### Distinct regional gene expression patterns are associated with *C. jejuni* and *Enterobacteriaceae* abundance in colonic tissues

Our data showed tissue-specific enrichment of *C. jejuni* and *Enterobacteriaceae* in C13+ *C. jejuni* mice (Fig. 5A-B). Therefore, we performed region-specific transcriptomic analysis to determine gene expression profiles for those specific regions. Our region of interest (ROI) was selected based on the presence (+) or absence (-) of *C. jejuni* (Fig. 7A). We observed an increased expression of cytokines (*Il17a, Il1a*, *Il22*), antimicrobial protein in Neutrophils (*Ngp*), epithelial repair and antimicrobial defense (*Saa3*) and immune modulation (*Marco*) in *C. jejuni* (+) region, whereas genes related to barrier function and microbial regulation (*Lypd8*, *S100a14*, *Muc13*) were predominantly expressed in *C. jejuni* (-) regions. These spatial expression patterns indicate that the mid colon mounts distinct transcriptomic responses depending on local bacterial abundance, with barrier and defense genes enriched in *C. jejuni* (-) zones (Fig. 7B, Additional file 5: https://figshare.com/s/b84ac4a3c99882822195). Strikingly, pathway enrichment analysis using Qiagen IPA revealed a significant upregulation of the atherosclerosis signaling pathway (Fig. 7C). This pathway encompasses key genes such as *Col3a1* (collagen type III alpha 1), *Ifng* (interferon gamma), *Il1a* and *Il1b* (interleukin 1 alpha and beta), *Itgb2* (integrin beta 2), *Lyz1/Lyz2* (lysozyme 1 and 2), *Pla2g7* (phospholipase A2 group VII), and *S100a8*. These genes are associated with extracellular matrix remodeling, immune cell recruitment, cytokine-driven inflammation, and dysregulation of lipid metabolism.

**Figure 7.**
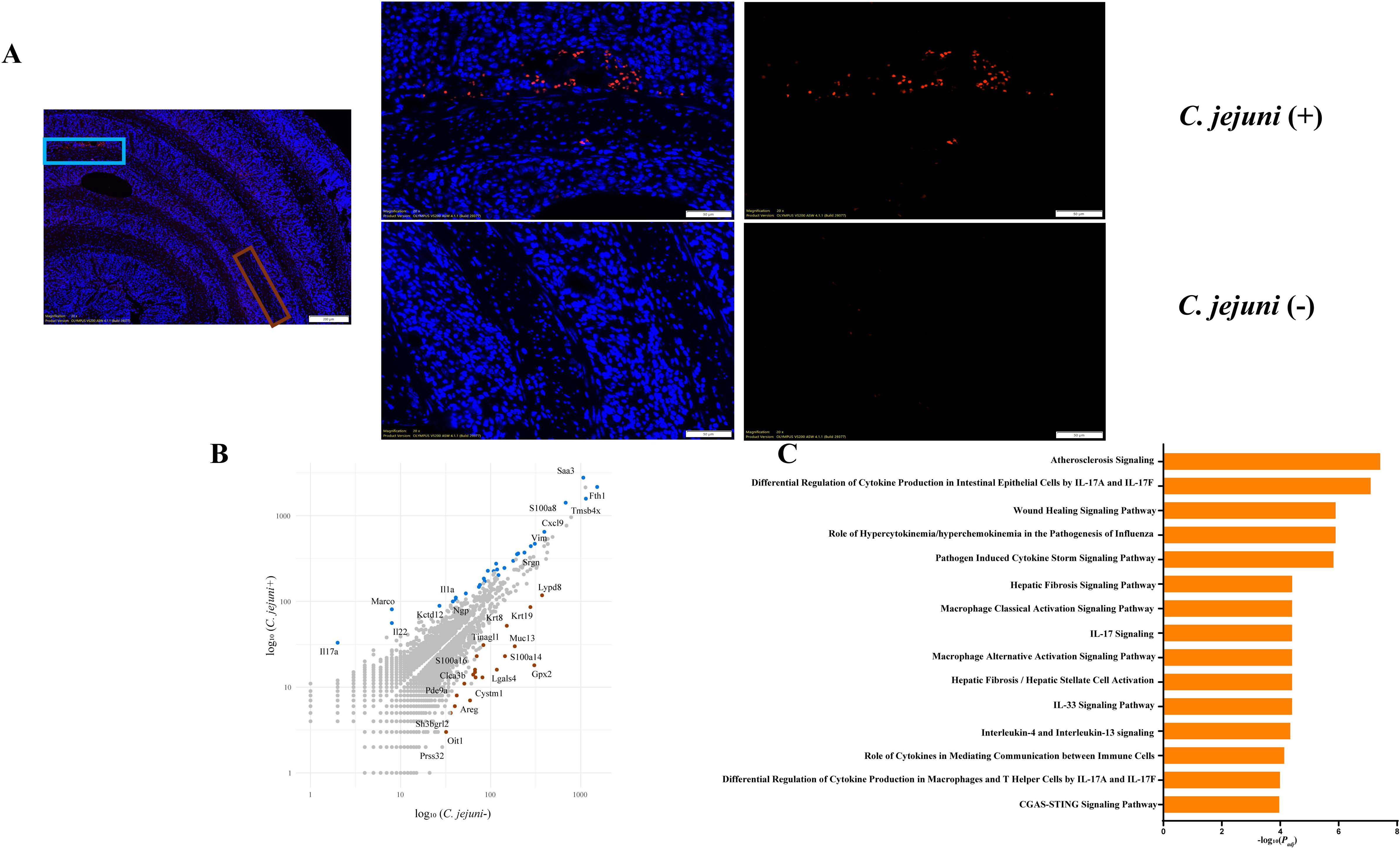
Region specific *C. jejuni* abundance reveals distinct transcription landscape (A) Detection of *C. jejuni* as represented by red dots in colonic sections of colonized mice were detected using FISH. Two ROIs were selected based on presence (+) or absence (-) of *C. jejuni* abundance. (B) Scatter plot showing significant differential expressed genes in *C. jejuni* (+) vs *C. jejuni* (-). Each dot represents a gene, gray: *P*_adj_ ≥ 0.05; light blue: increased expression in *C. jejuni* (+) and *P*_adj_ < 0.05; brown: increased expression in *C. jejuni* (-) and *P*_adj_ < 0.05. (C) Top host pathways significantly upregulated in *C. jejuni* (+) region as compared to *C. jejuni* (-) region in *C13 + C. jejuni* infected mice. Qiagen IPA software was used for this analysis (*P*_adj_ value < 0.05).

RNAscope analysis of the *Enterobacteriaceae* revealed regional differences in signal intensity across the C13 + *C. jejuni* colon. Spatial analysis on *Enterobacteriaceae* high region exhibited elevated expression of genes (*Ngp*, *Ila*, *Ilb*, *Ubd*, *S100a8*, *S100a9* and *Nos2*) involved in host defense, pathogen recognition, and cellular stress, compared to *Enterobacteriaceae* low regions (Fig 8 A-C, Additional file 6: https://figshare.com/s/b84ac4a3c99882822195). Common inflammatory pathways between *C. jejuni* and *Enterobacteriaceae* ROI were significantly enriched, including the *Il-17a* and *Il-17f*-mediated differential regulation of cytokine production in both intestinal epithelial cells and macrophages, and the pathogen-induced cytokine storm signaling pathway, indicating epithelial and systemic immune activation. Furthermore, both the classical and alternative activation signaling pathways of macrophages were upregulated. The alternative activation pathway was characterized by increased expression of genes such as *Cebpb*, *Chil3/Chil4*, *Dusp1*, *Il1a*, *Il1b*, and *Marco*, while the classical activation pathway featured elevated levels of *Ccl5*, *Cxcl9*, *Ifng*, *Il17a*, *Il1a*, and *Il1b*. Unique pathways include activation of the cGAS– STING pathway and cytokine signaling by immune cells in *C. jejuni* high regions, reflecting intracellular pathogen sensing and immune modulation. In contrast, *Enterobacteriaceae* high areas exhibited upregulation of antimicrobial peptides and Toll-like receptor (TLR) signaling pathways, indicative of classical mucosal defense mechanisms against extracellular bacteria. These findings, together with the upregulation of signaling pathways, reflect a complex and diverse spectrum of macrophage activation states within the tissue (Fig. 7C).

**Figure 8.**
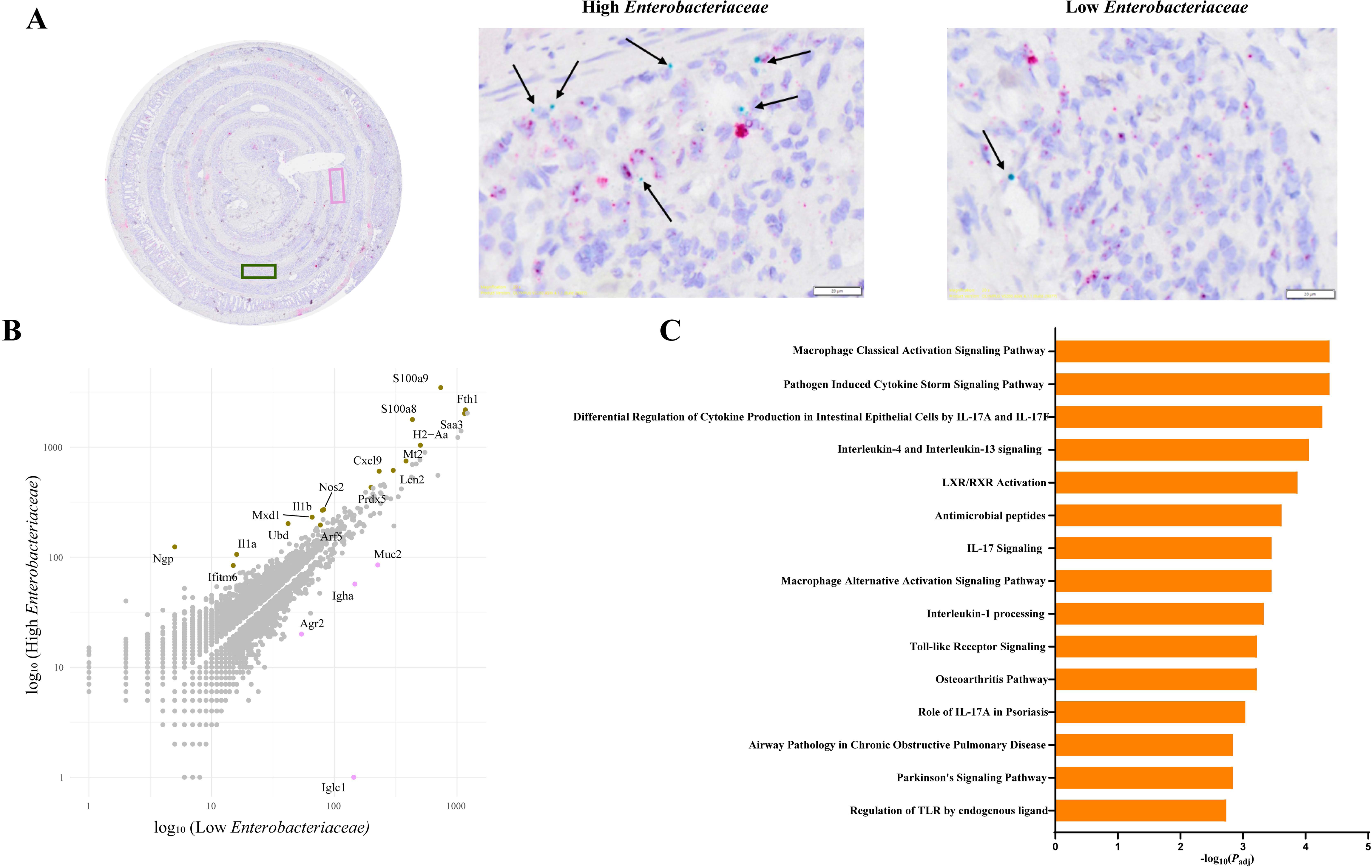
Region specific *Enterobacteriaceae* abundance reveals distinct transcription landscape (A) Detection of *Enterobacteriaceae* as represented by green dots in colonic sections of colonized mice were detected using RNAscope. Two ROIs were selected based on high or low *Enterobacteriaceae* abundance. (B) Scatter plot showing significant differential expressed genes in High *Enterobacteriaceae* vs Low *Enterobacteriaceae*. Each dot represents a gene, gray: *P*_adj_ ≥ 0.05; green: increased expression in High *Enterobacteriaceae* and *P*_adj_ < 0.05; pink: increased expression in Low *Enterobacteriaceae* and *P*_adj_ < 0.05. (C) Top host pathways significantly upregulated in High *Enterobacteriaceae* region as compared to Low *Enterobacteriaceae* region in *C13 + C. jejuni* infected mice. Qiagen IPA software was used for this analysis (*P*_adj_ value<0.05).

Using spatial transcriptomic data, we performed cell-type deconvolution to examine differences in cellular composition across GF, C13, and C13 + *C. jejuni* mice (Supplemental Fig. S2 A–C). In both GF and C13 groups, we observed broad expressions of genes associated with enterocytes and goblet cells, indicating preservation of the epithelial compartment. In contrast, a substantial loss of these epithelial and goblet cell signatures was observed throughout the colon of C13 + *C. jejuni* group. The most striking change was a notable increase in genes associated with myeloid cell populations, reflecting innate immune response as previously reported (35). These findings suggest that *C. jejuni* colonization drives a major remodeling of the cellular landscape in the colon.

## Discussion

In this study, we present a novel tractable bacterial consortium (C13) to study *C. jejuni* host response during inflammatory response. We observed that the C13 consortium alone does not promote an acute inflammatory response in colitis-susceptible *Il10^-/-^* mice, suggesting the absence of inflammatory drivers within the consortium. The addition of *C. jejuni* to the consortium significantly triggered colitis, suggesting that this pathogen can alter host-bacterial interaction. These host responses replicate the hallmarks of campylobacteriosis, such as increased inflammatory gene transcripts, neutrophil infiltration, and *C. jejuni* invasion (11,35). A recent study reported the use of a bacterial consortium, OligoMM^12^ to study host response to various enteropathogenic bacteria, including *C. jejuni* (55). Although our RNA-seq was performed on fecal samples, several transcriptional changes overlapped with genes reported as highly upregulated or downregulated in the cecal transcriptome of the OligoMM^12^ model (55), underscoring the biological relevance of our findings. However, the OligoMM^12^ model elicits only mild cecal intestinal inflammation with limited hallmarks related to Campylobacteriosis.

In our model, we noted an increased abundance of two commensals based on the consortium status: *P. distasonis* in C13 group and *P. bifermentans* in the C13 + *C*. *jejuni* group. While *P. distasonis* plays a critical role in maintaining gut barrier integrity by increasing the frequency of ILC3 cells (56), *P. bifermentans* has been shown to increase inflammatory cytokine levels and damage intestinal epithelial cells in a DSS acute colitis model. Whether *P. distasonis* synergizes with *C. jejuni* to promote intestinal inflammation is unknown and will require modification of the C13 composition to address its specific role. Interestingly, we observed an increased abundance of pks^+^ *E. coli* in the stool samples of C13 + *C. jejuni* mice. A similar increased abundance of *Enterobacteriaceae* was reported in mice infected with *C. jejuni* (43). Intestinal inflammation has been shown to foster the expansion of facultative anaerobic bacteria such as *E. coli* by impairing epithelial cells’ ability to perform β-oxidation and oxidative phosphorylation, a process resulting in higher oxygen tension in the gut (57). Another study revealed that host-derived nitrates produced during inflammation increased the abundance of *E. coli* that utilize nitrates for respiration (58).

Whether *C. jejuni* fosters *E. coli* growth through similar mechanisms will require further investigation. Since *P. bifermentans* is a strict anaerobic bacterium, other factors are likely in play regarding reshaping the abundance of this bacterium. Our fecal RNA-seq analysis showed that the *Slc7a11* transporter which is involved in the uptake of extracellular cysteine (46) was significantly up-regulated in C13 + *C. jejuni* compared to C13. *C. jejuni* has also been shown to utilize host-derived cysteine for its metabolism, as it lacks a *de novo* mechanism for cysteine production (59). *E. coli* metabolizes host-derived cysteine for the production of hydrogen sulfide which plays a protective role against oxidative stress (60). This could suggest a potential mechanism by which *C. jejuni* and *E. coli* adapt to the changing intestinal micro- environment.

We also observed differential expression of genes associated with bacterial virulence such as PorA (*C. jejuni*), Dps (*C. jejuni*), aspartate ammonia lyase (*C. jejuni* and *E. coli*), peroxiredoxin (*C. jejuni* and *P. bifermentans*) when *C. jejuni* is added to C13, which suggests an adaptive and protective mechanism to survive in the gut. We evaluated virulent genes involved in biofilm, defense, motility, stress response and quorum sensing in *E. coli*. We observed increased BasR expression which encodes for the two-component regulatory system BasS/BasR (defense), FliA that encodes for flagellar expression (motility), Qseb which is a part of quorum sensing regulatory cascade, RpoE encoding for a sigma factor that regulates stress response, and AraC influencing biofilm formation. This increase in *E. coli* virulence gene expression may reflect potential interaction with *C. jejuni* and host-derived inflammation. Interestingly, our findings align with a previous study demonstrating that *C. jejuni* activates latent virulence gene expression in *E. coli* that subsequently induces pro-inflammatory gene expression (61).

Spatial transcriptomic analysis revealed distinct host expression patterns between GF and C13-colonized mice. When compared to C13-colonized mice, GF animals exhibited elevated expressions of genes such as *Cyp2d10* and *Ighg1*. *Cyp2d10*, a member of the cytochrome P450 family, involved in xenobiotic metabolism, suggesting increased cellular stress or metabolic adaptation without microbial colonization. In contrast, expression of these genes was significantly dampened in C13-colonized mice, suggesting that even a simplified microbial consortium can attenuate immune activation by promoting immune tolerance.

Moreover, C13-colonized mice exhibited increased expression of key epithelial defense genes including *Reg3g*, *Muc2*, and *Ang4*. These findings suggest that the C13 consortium enhances basal epithelial defense by upregulating key barrier and antimicrobial genes, *Muc2*, *Reg3g*, and *Ang4*, as compared to GF mice, potentially priming the gut mucosa for greater resilience against pathogenic insults. Interestingly, both GF vs C13 + *C. jejuni* and C13 vs C13 + *C. jejuni* comparisons showed upregulation of genes related to inflammation and innate immune activation, including *Saa3*, *S100a8*, *S100a9*, *Lcn2*, *Chil3*, and *Ngp*. Conversely, genes upregulated in GF or C13 alone, such as *Slc26a3*, *Sycn*, *Tgm3*, and *Gsdmc3*, were associated with epithelial barrier maintenance, electrolyte transport, and digestive function. These findings indicate that *C. jejuni* infection drives a shift from homeostatic epithelial gene expression toward an inflammatory state, regardless of baseline microbiota status. Additionally, the observed downregulation of pathogen induced cytokine storm pathway in Cluster 17 contrasts with their upregulation in other clusters, suggesting region-specific modulation of inflammation related signaling. Given that Cluster 17 is localized predominantly to the distal colon, this may reflect a localized protective or stress-response environment that suppresses pro-inflammatory response to *C. jejuni* infection. In contrast, the higher abundance of *C. jejuni* in the mid colon, corresponding with Clusters 2 and 3, aligns with increased activation of pro-inflammatory and immune-modulatory pathways. Notably, in the C13 + *C. jejuni* infected mice, increased levels of *E. coli* and *P. bifermentans* were also observed, which may contribute to the complex microbial interactions influencing inflammation signaling. These spatial differences also imply that host may exert differential effects on the colonic microenvironment depending on regional *C. jejuni* colonization, promoting immune and inflammatory signaling in some areas while triggering protective responses in others. Further studies are needed to elucidate the host derived mechanisms driving this regional specificity and to determine whether these findings translate into differences in tumor development or progression. Our cell-type deconvolution analysis revealed a pronounced spatial enrichment of myeloid cells and depletion of enterocytes and goblet cells within the colon of C13 + *C. jejuni*-infected mice. This suggests that *C. jejuni* infection in the presence of the defined microbiota triggers a localized host immune response geared toward pathogen clearance and tissue remodeling.

Region-specific spatial profiling revealed that host responses to bacterial stimuli are highly localized and shaped by the composition of adjacent microbial communities. In regions enriched with *C. jejuni*, we observed signatures of intracellular pathogen recognition *via* the cGAS–STING pathway and elevated cytokine responses. These responses were accompanied by increased expression of *Vim* and *Marco*, suggesting that *C. jejuni* not only triggers inflammatory signaling but may also induce epithelial stress and activate tumor-associated macrophage programs. This gene repertoire pattern aligns well with our findings that *C. jejuni* carrying the genotoxin cytolethal-distending toxin (CDT) promotes the development of primary and metastatic CRC (9,10). In contrast, *Enterobacteriaceae-*dominated regions exhibited robust expression of antimicrobial peptides and TLR-related genes, consistent with classical mucosal defense mechanisms. Together, these spatially restricted host responses underscore how specific bacterial taxa, even within a defined microbial consortium, elicit divergent immune and epithelial programs that may influence inflammation and disease susceptibility. Overall, our findings emphasize the value of combining gnotobiotic models with high-resolution spatial and molecular tools to dissect the complex dynamics between the host, microbiota, and pathogens. This approach not only advances our understanding of microbial influence on intestinal immunity but also opens new avenues to design targeted microbial therapies that leverage defined consortia for disease prevention or treatment.

### Limitations of this study

While this study provides valuable insights into host–microbe and pathogen interactions using spatial transcriptomics and gnotobiotic models, several limitations should be acknowledged. Spatial transcriptomic analysis offers a static snapshot of gene expression, limiting our ability to capture dynamic changes in microbial localization and host responses over time. Additionally, the resolution of current spatial transcriptomics technology and the sequencing depth achieved may not fully capture cellular heterogeneity or detect low-abundance transcripts, potentially overlooking rare but important cell populations or subtle transcriptional changes. Although bacterial transcriptional programs were modulated during infection, the potential role of host-derived factors in driving these changes remains speculative and requires further investigation. Notably, whether *C. jejuni* itself can alter host-derived signals that in turn regulate microbial behavior is an exciting question that warrants further study.

## Conclusion

In conclusion, our study establishes a bacterial consortium allowing the study of complex host responses to *C. jejuni* infection. Our model replicates all the major classic campylobacteriosis hallmarks, such as increased inflammatory gene transcripts in the host and neutrophil infiltration at the colon sites following *C. jejuni* infection. Spatial transcriptomic profiling of the entire colon further revealed that host responses to microbial stimuli are highly compartmentalized, emphasizing the critical role of local tissue microenvironments in shaping epithelial and immune dynamics.

## Acknowledgments

The authors thank the University of Florida Animal Care Services—Germ-free Division for their assistance with germ-free husbandry. We are grateful to Dr. Rachel Newsome for her assistance with the IsoPositive Cage System. We would also like to thank the UF Molecular Pathology Core for the H&E staining service. UF-Health Cancer Center ICBR cores, and for their support in Qiagen IPA software, bacterial RNA-sequencing, and spatial transcriptomics. Graphical abstract was created using Biorender.com and is licensed for publication.

## Declarations

### Ethics approval and consent to participate

All animal experiments were approved by the Institutional Animal Care and Use Committee (IACUC) at the University of Florida (UF) and performed at UF Animal Care Facilities (IACUC Protocol #202200000637).

### Consent for publication

Not Applicable

### Availability of data and materials

Sequencing reads have been deposited in the National Center for Biotechnology Information (NCBI) Sequence Read Archive (SRA) under bioproject ID: PRJNA1209065 (16S rRNA), PRJNA1209059 (RNA-seq). Spatial transcriptomics data have been deposited in the NCBI Gene Expression Omnibus (GEO) under accession ID: GSE300781.

### Authors contributions

Conception: SC, R.Z.G, CJ. Methodology: SC. Investigation: SC, R.Z.G, CJ. Manuscript drafting/revisions: SC, R.Z.G, CJ. Bioinformatic analysis: R.Z.G. Supervision: CJ. All authors read and provided final approval of the submitted manuscript.

---

**Supplementary Figure 1.**
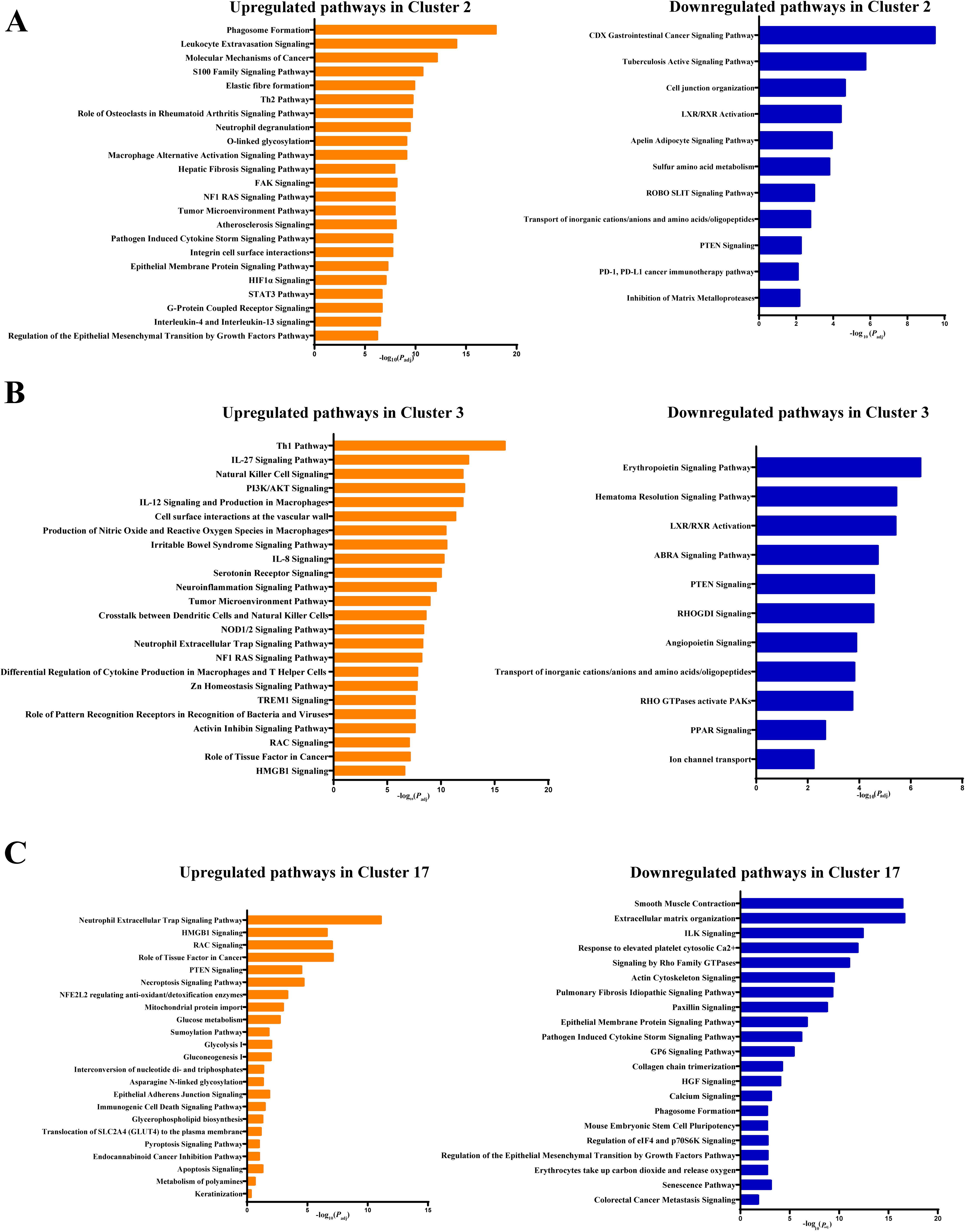
Pathway analysis of Clusters 2,3, and 17 differentially expressed genes in the host. (A) Upregulated pathways in Cluster 2, Downregulated pathways in Cluster 2, (B) Upregulated pathways in Cluster 3, Downregulated pathways in Cluster 3, (C) Upregulated pathways in Cluster 17, Downregulated pathways in Cluster 17. Qiagen IPA software was used to identify significantly altered pathways (*P*_adj_ value < 0.05).

**Supplementary Figure 2.**
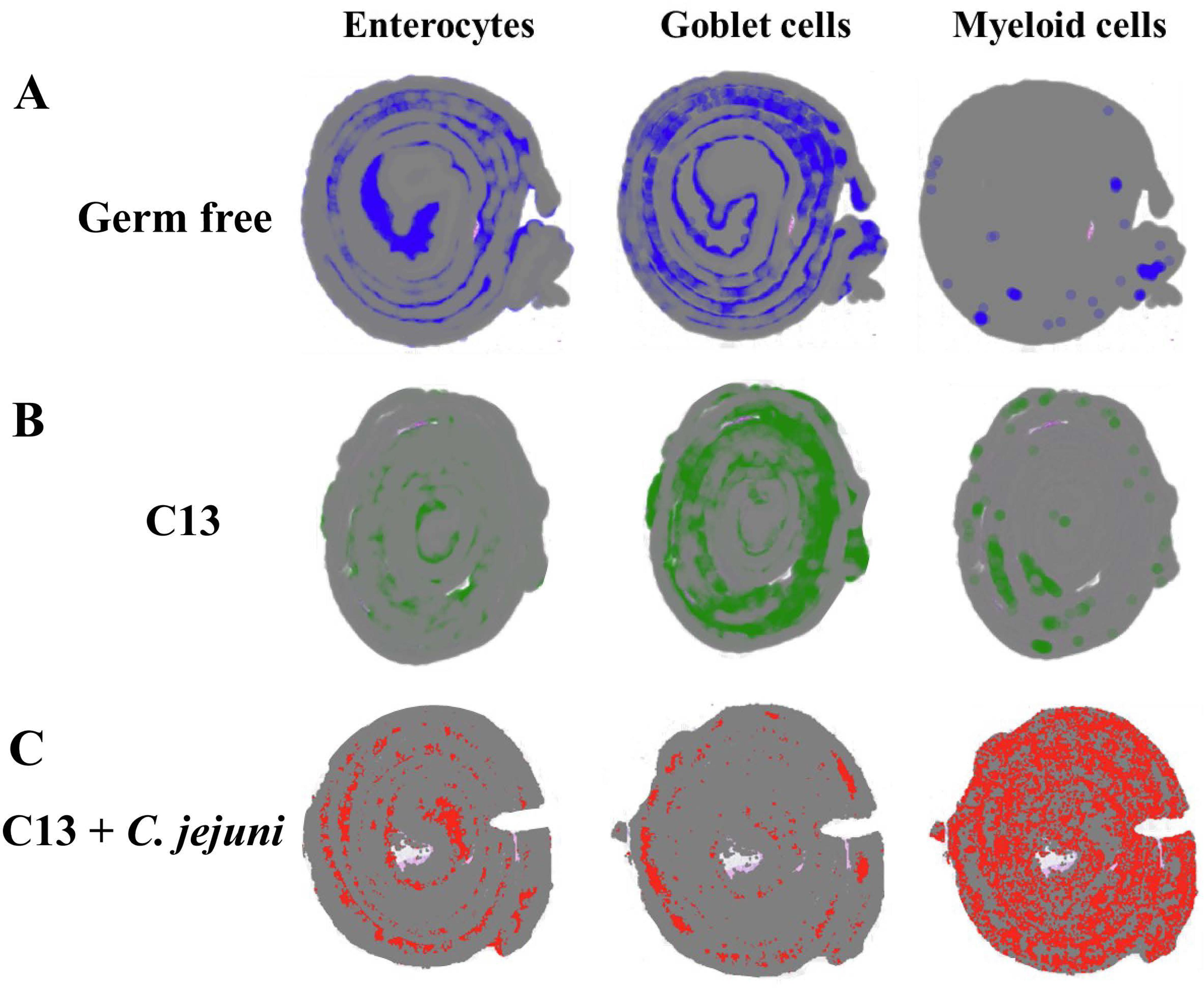
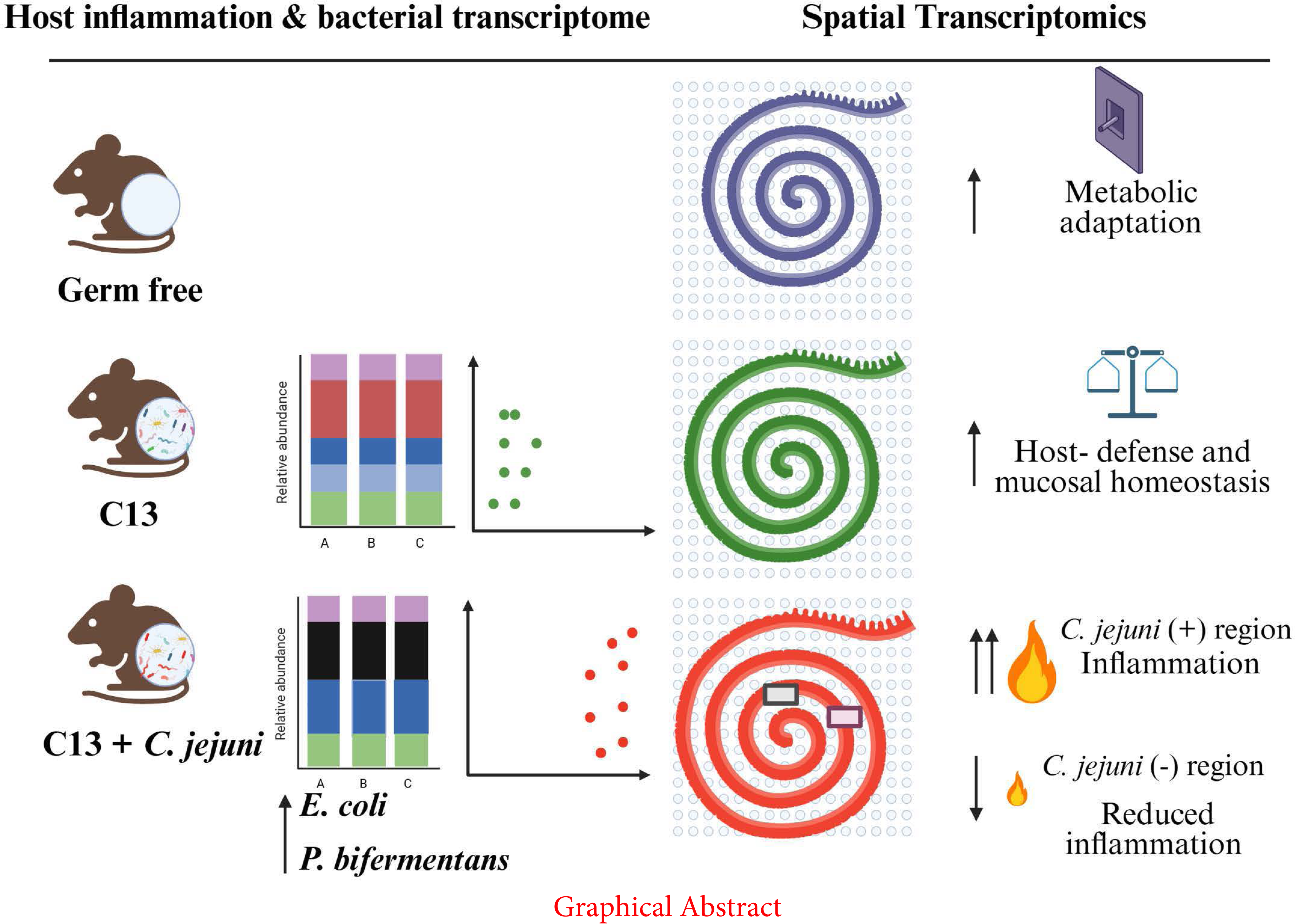
*C. jejuni* infection drives spatial enrichment of myeloid cells in the colon (A– C) Spatial cell-type deconvolution of major cell types in colonic Swiss roll sections from (A) germ-free (GF), (B) C13-colonized, and (C) C13 + *C. jejuni*-infected mice.

Additional files available on Figshare at https://figshare.com/s/b84ac4a3c99882822195:

**Additional file 1**: Mouse RNA-seq data table listing gene symbols, product, LogFC, P-value, FDR and significance.

**Additional file 2**: Bacterial RNA-seq data table listing gene IDs, product, LogFC, P-value, FDR and significance.

**Additional file 3**: Spatial transcriptomics clusters 2, 3 and 17 lists, showing up and down regulated genes. Related to Figure 6 A and B.

**Additional file 4**: Differentially expressed genes between GF vs. C13, GF vs. C13CJ and C13 vs. C13CJ. Related to Figure 6C.

**Additional file 5**: Significant differentially expressed genes between *C. jejuni* (+) vs. *C. jejuni* (-). Related to Figure 7B.

**Additional file 6**: Significant differentially expressed genes between High *Enterobacteriaceae* vs. Low *Enterobacteriaceae.* Related to Figure 8B.

